# Enhancer Placement Impacts Transcriptional Dynamics in *Drosophila* Embryos

**DOI:** 10.1101/2025.08.14.670187

**Authors:** Emilia A. Leyes Porello, Jiayi Wu, Samantha Fallacaro, Emma Dreispiel Juan, Robert T. Trudeau, Jose Andres Vidal, César Nieto, Abhyudai Singh, Mustafa Mir, Bomyi Lim

## Abstract

The functional role of enhancer-promoter configurations in transcription regulation remains poorly understood, despite the wide range of linear genomic distance and relative enhancer positioning observed in endogenous contexts. While canonical models suggest that enhancers operate independently of genomic position, technical limitations have obscured insights on transcriptional kinetics. Here, we employ single-cell MS2/MCP-based live imaging in *Drosophila* embryos to systematically analyze transcriptional dynamics driven by sub-10kb enhancer-promoter arrangements. Kinetic analyses reveal that while linear enhancer-promoter distance moderately tunes transcriptional output, downstream enhancer positioning reduces mRNA output by 70%. Each configuration modulates distinct transcriptional parameters: linear distance governs initiation kinetics, while relative enhancer positioning dictates transcriptional stability. These effects are consistent across varied enhancer and reporter sequences, revealing configuration-dependent *cis-*regulatory element arrangement as an intrinsic mechanism for transcriptional fine-tuning. This work challenges the paradigm of configuration-independent enhancer function and establishes a framework to dissect the interplay between genome architecture and *trans-*acting factors.

## Introduction

The complex nature of transcription regulation continues to challenge our complete understanding of the multilayered regulatory logic that governs gene expression programs critical for organismal development. While extensive research has established core principles of transcriptional control, such as enhancer-mediated recruitment of transcriptional machinery and subsequent promoter activation, the dynamic interplay between diverse *cis*-regulatory elements presents a multifaceted regulatory landscape. Epigenetic regulators, such as boundary elements and chromatin remodelers, add further layers of mechanistic complexity by modulating the proximity of cognate elements to one another, which is implicated in the control of transcriptional output ^1–3^. Further intricacy is granted by *trans-*acting elements like transcription factors (TFs) and their higher order assembly into hubs or condensates, complicating efforts to delineate the underlying physical mechanisms governing gene regulation. This intricate web of regulatory interactions underscores the ongoing challenge in fully understanding the nuanced and dynamic nature of gene expression control in biological systems.

Even in controlled reporter settings where a single enhancer regulates a cognate gene, the underlying regulatory principles are not fully understood. In 1981, Banerji et al. characterized the first regulatory enhancer in the simian virus 40 (SV40) genome, by identifying a short DNA segment that can dramatically boost transcription from a target promoter ^4^. In the decades following this groundbreaking discovery, the role of enhancers in regulating gene expression has been extensively probed. However, two key assumptions have persisted largely unchallenged: that enhancer orientation (sense vs. antisense) and positioning relative to target promoters (upstream vs. downstream) have insignificant bearing on enhancer functionality ^4–8^.

Enhancers exhibit remarkable flexibility in their genomic positioning, executing their regulatory functions despite being endogenously located at varying distances from their target genes – ranging from promoter-proximal sites to regions hundreds of kilobases away – and at varied relative positions, with many endogenous genes regulated by both upstream and downstream enhancers ^9^. For instance, the *even-skipped* (*eve*) gene in *Drosophila* relies on an array of stripe-specific enhancers positioned on both sides of the gene, each playing a distinct, yet critical, role in tightly controlling its expression ^10^. Differences in transcription factor recruitment are known to underlie these divergent roles, though any influence of relative enhancer-promoter (E-P) placement in this regulatory context is uncharacterized. While higher-order chromatin organization likely facilitates long-range enhancer-promoter interactions in endogenous settings, early studies using reporter systems claimed that enhancers exhibit positional and orientational independence even at smaller length scales ^4,5^. In their seminal experiment, Banerji et al. showed that inserting the 72 bp SV40 enhancer either 1400 bp upstream or 3300 bp downstream of the rabbit beta-globin gene promoter – regardless of orientation – successfully activated transcription in HeLa cells ^4^. The immunoglobulin G (IgG) heavy chain enhancer was also described as both positioning and orientation independent – capable of activating tissue-specific transcription when inverted or moved downstream of the promoter ^5^. While groundbreaking, these studies could not detect quantitative expression differences driven by enhancers in varied genomic contexts, due to technical limitations that prevented high spatiotemporal resolution and efficient genome editing. More advanced, quantitative techniques in imaging and genomic analysis now enable a more detailed characterization of these enhancer behaviors and the kinetics of the transcriptional activity they regulate.

Recent advancements in genome-wide techniques, such as chromatin conformation capture (3C) and its derivatives, along with single-cell, fixed imaging methods like fluorescent *in-situ* hybridization (FISH), have significantly enhanced our understanding of gene regulation ^11–13^. However, these approaches have inherent limitations that either obscure individual cell behavior due to bulk-averaged data (in genome-wide studies) or lack temporal resolution as a result of tissue fixation (in single-cell, fixed imaging assays). Further, most attempts to understand the mechanisms of E-P interactions have been done at long-range genomic scales, which are largely governed by boundary or tethering elements and other epigenetic regulatory mechanisms ^14–16^. To elucidate the fundamental regulatory principles of enhancer elements, it is crucial to investigate the mechanisms of direct enhancer-mediated transcriptional regulation at shorter distances, distinguishing them from effects that dominate at larger length scales.

In this study, we designed reporter constructs to systematically vary enhancer-promoter configurations in early *Drosophila* embryos and quantified single-nucleus transcriptional activity in real-time by employing the MS2/MCP system. A striking result emerged when mRNA output driven by downstream enhancer placement was reduced by 70% compared to an equidistant, upstream enhancer. To probe the underlying regulatory grammar, we conceptualized transcriptional activation as a two-part phenomenon: first, the initial, post-mitotic formation of a productive E-P interaction initiating transcriptional activity. Second, the stability of the active state throughout the remainder of the cell cycle. Our data suggest that, at sub-10 kb lengths scales, linear E-P distance primarily influences transcriptional onset time with increasing genomic distance, delaying activation and mildly inversely correlating to transcriptional output levels. Further, we report that relative enhancer positioning modulates active state stability with downstream enhancer placement driving significantly less stable transcription and yielding drastically reduced mRNA production compared to upstream enhancer counterparts.

These findings not only provide novel insights to enhancer-mediated transcriptional regulation but also call for a fundamental revaluation of our understanding of enhancer function. By challenging the long-held notion of configuration-independent activity, our research paves the way for a more comprehensive model of gene regulation.

## Results

### Relative enhancer positioning and enhancer-promoter distance finely tune transcriptional activity

To investigate the role of E-P configuration in transcription regulation, we developed a series of seven reporter constructs with systematic variations in E-P distance and relative enhancer positioning. These constructs ranged in linear E-P distance from 0 kb to 10 kb, with five featuring enhancers upstream of their target promoter (EnhUP) and two with enhancers placed downstream (EnhDOWN). All constructs shared identical regulatory element sequences and were site-specifically inserted to a low-complexity region on chromosome III of *Drosophila melanogaster* ^17^. This strategic placement allowed us to isolate the effects of enhancer-promoter configurations on transcriptional behavior while minimizing confounding effects from the surrounding chromatin environment since each configuration operates within the same genomic background. *Drosophila* provides an ideal model system for this systematic experimental design, as the median distance between enhancers and their target promoters is approximately 10 kb, offering a biologically relevant context to explore regulatory mechanisms that prevail at these distances ^9^. Moreover, systematic analysis at this short length-scale enables investigation of the direct effect of E-P interactions on transcription, without indirect effects from auxiliary factors like insulators and tethering elements.

The constructs each contained a *lacZ* reporter gene driven by the *snail* distal enhancer (snaDE) and the minimal 100-bp *even-skipped* promoter (evePr), a combination known to yield strong expression in early *Drosophila* embryos ^18^. The snaDE, endogenously located ∼7 kb upstream of the *snail* promoter, drives robust ventral mesodermal expression during early *Drosophila* embryogenesis ^19^. Its activity depends on synergistic recruitment of Dorsal (Dl), Twist (Twi), and Zelda (Zld) transcription factors (TFs) through clustered low-affinity binding sites ^20^. This cooperative regulation establishes a sharp dorsoventral expression boundary critical for mesoderm invagination at the end of nuclear cycle 14 (NC14).

We designed an array of constructs with systematic variations in E-P distance by inserting neutral spacer sequences between snaDE and evePr. These neutral spacers were designed to lack binding sites for the three key TFs regulating snaDE in the early *Drosophila* embryo (Dl, Twi, and Zld). We identified a 5 kb fragment of the *E. coli* gene *yeeJ* (Figure 1A) containing only a single Zld binding motif, and mutated the site to use the gene fragment as a spacer sequence. We additionally utilized a published, synthetic spacer sequence, computationally designed to lack binding sites for all active TFs in early *Drosophila* embryos ^21,22^. These modular spacer blocks were then assembled in various lengths, enabling precise control of the linear spatial relationship between enhancer and promoter elements, while avoiding the introduction of putative TF binding sites.

**Figure 1.**
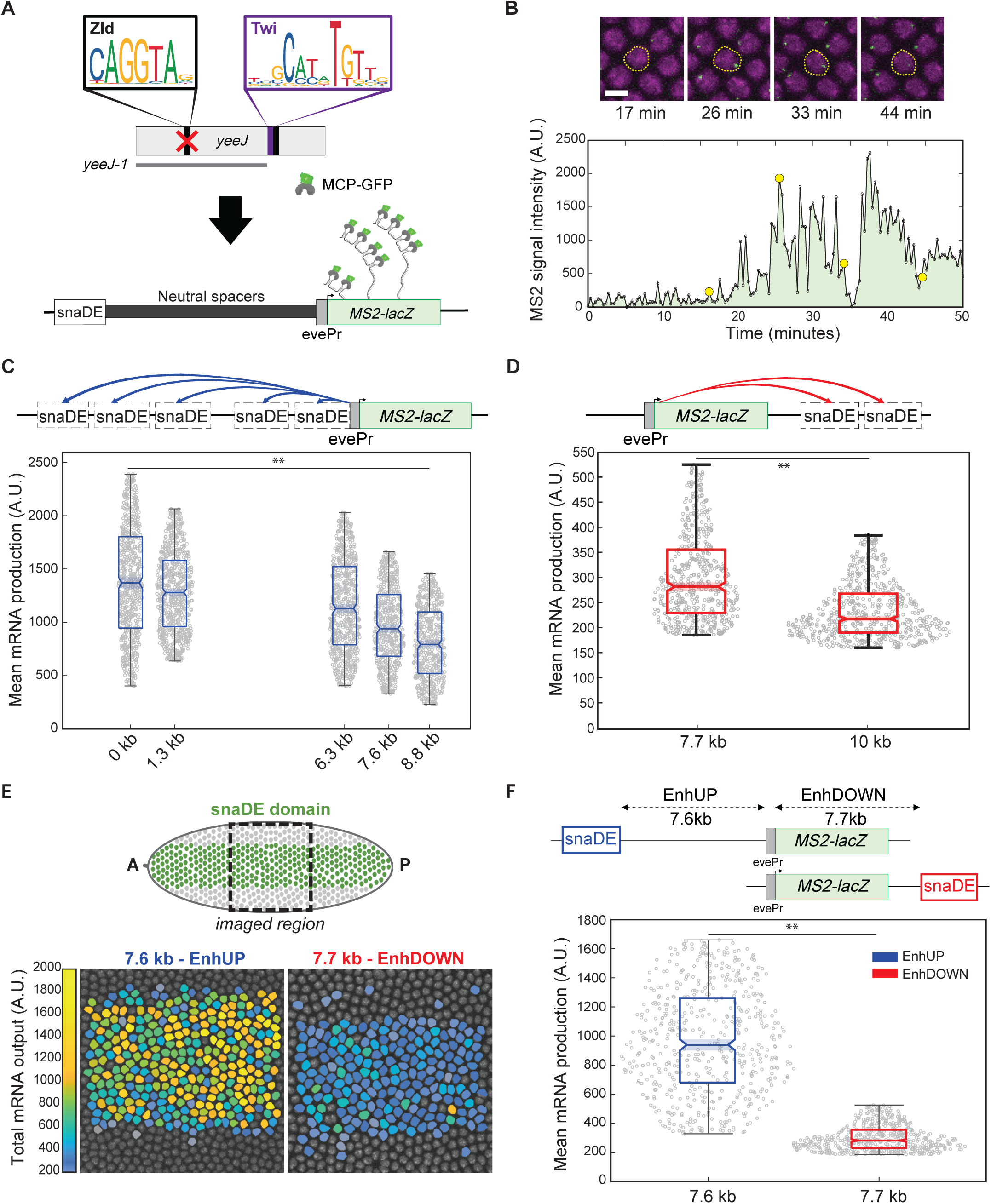
Relative enhancer positioning and enhancer-promoter distance finely tune transcriptional activity. (A) Schematic of neutral spacer designed by deletion of Zld binding motif in *E. coli* gene, *yeeJ*. (B) Representative MS2 trajectory and corresponding snapshots (marked in yellow circles) for a single nucleus over time in NC14. The area under the trajectory can be used as a proxy for mRNA production. (C,D,F) Mean mRNA produced by all active nuclei in NC14 for (C) EnhUP, (D) EnhDOWN, and (F) mid-distance, opposing enhancer positions. Dots represent total mRNA produced by individual active nuclei over the full NC14. Each data set contains combined data for 3 biological replicates. Number of nuclei in each data set as follows: n(0kb)=922, n(1.3kb)=915, n(6.3kb)=888, n(7.6kb)=991, n(7.7kb)=681, n(8.8kb)=840, n(10kb)=809. Statistical significance determined by ANOVA for C and student’s t-test for D and F with α=0.05 for both. ** denotes p<0.01. (E) Schematic of the snaDE expression domain (top) and imaged region along with false colored embryo snapshot depicting total mRNA produced by each nucleus in the pictured embryo (bottom). EnhDOWN construct drives lower mRNA production uniformly across the *snail* expression domain.

We employed the bacteriophage-derived MS2/MCP live imaging system to quantify transcriptional activity driven by these varied E-P configurations ^23–25^. The 24x MS2 sequence was inserted in the 5’UTR of the *lacZ* reporter. Upon transcription, this sequence generates 24 RNA stem loops with high affinity for the GFP-tagged MS2 coat protein (MCP) (Figure 1A). As the MCP-GFP fusion proteins are recruited to these stem loops at the transcription site, a fluorescent signal is produced, providing a real-time measure of instantaneous transcriptional activity in living cells (Figure 1B). We first confirmed neutrality of the spacer sequences by expressing a control construct lacking an enhancer but containing all spacer and reporter fragments. Only a few nuclei in the entire *sna* expression domain exhibited transient and weak transcription activity, presumably driven by the evePr (Supplemental Figure 1).

To analyze the impact of E-P distance on transcription, we compared the amount of mRNA produced from each construct. The cumulative transcriptional output for each construct was quantified by integrating the MS2 fluorescence intensity of each transcriptionally active nucleus over the full duration of NC14, spanning from mitosis to the onset of gastrulation (Figure 1B). Analysis across all configurations revealed a modest, non-linear inverse correlation between E-P distance and mean mRNA produced per nucleus, with mRNA output falling as distance increased (Figures 1C and 1D). This trend is consistent with that observed by Yokoshi et al. in *Drosophila* embryos where increasing E-P distance from 6.5 kb to 9 kb yielded > 50% reduction in the total amount of RNA synthesis ^26^. Similar trends have also been reported at larger E-P distances (hundreds of kilobases) in mouse embryonic stem cells (mESC) ^2,3,27^.

While the distance-output correlation was observed regardless of enhancer placement upstream or downstream of the promoter, a more pronounced effect on mRNA production emerged when comparing across opposing enhancer positions. Notably, downstream enhancers (EnhDOWN) drove significantly reduced transcriptional output compared to their upstream (EnhUP) counterparts, even at nearly identical E-P distances (Figure 1F), suggesting a strong positional bias in enhancer function at this scale. This is in stark contrast to the classical notion that enhancers function in a positioning-independent fashion ^4–8^.

In light of this striking result, we sought to understand the coordinated, embryo-scale behavior leading to such reduced transcriptional output in the downstream enhancer configurations. When visualizing the total mRNA produced per nucleus of the embryos for mid-range (7.6kb and 7.7kb) constructs with opposing enhancer positioning, we find that transcriptional output is uniformly regulated across the snaDE expression domain for both constructs (Figure 1E). Further, the boundary width of the expression domain and the number of active nuclei within the expression domain are not significantly affected by E-P configuration (Supplemental Figure 2). This suggests that the transcriptional kinetics of actively transcribing nuclei are uniformly influenced by enhancer position and that these give rise to differences in mRNA output. We thus turned to detailed kinetic analysis of the transcriptional trajectories produced by each construct to shed light on the biological mechanisms underlying these E-P configuration-dependent trends in mRNA production.

### Enhancer-promoter configuration drives two distinct properties of transcription regulation: onset time and active state stability

To investigate the mechanisms underlying the observed differences in transcriptional output, we analyzed the collective transcriptional activity of all active nuclei during NC14 using heatmaps for various E-P configurations. These visualizations revealed that the rate of initial transcriptional activation decreases with increasing enhancer-promoter distance, as reflected by delays in the timing at which 50% of nuclei became active (t_50%_), independent of the relative enhancer positioning. Additionally, downstream enhancer configurations exhibited reduced signal amplitude and more stochastic transcriptional states compared to upstream configurations (Figure 2A).

**Figure 2.**
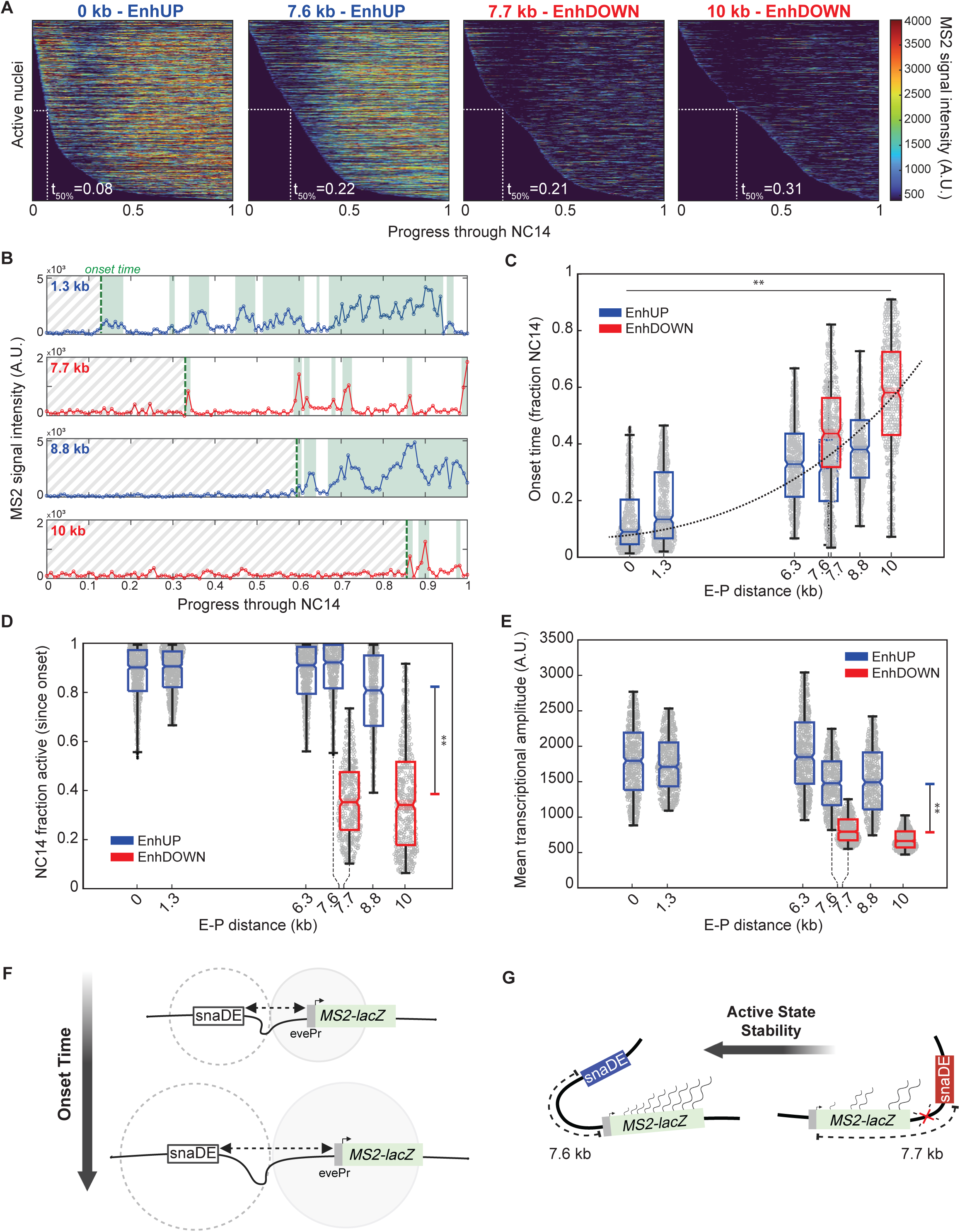
Enhancer-promoter configuration drives two distinct properties of transcription regulation: onset time and active state stability. (A) Heat map of transcriptional trajectories for all active nuclei in each construct. Each map contains combined data for three biological replicates. The timing of activation of half the total active nuclei is reported as t_50%_. (B) Representative MS2 trajectories for nuclei in each of four shown configurations over full NC14. Dashed green line represents the onset time. Shaded green region represents active transcription based on thresholding. Gray slashed fill represents time prior to onset time. (C) Onset time plotted for each shown configuration. Dashed line is a best-fit trend through the median value of each data set. (D) Fraction of NC14 after onset time that each nucleus spends in the active state plotted for each construct. (E) Mean transcriptional amplitude of the active state plotted for each shown configuration. Statistical significance between EnhUP and EnhDOWN subsets were performed by combining the data into two groups based on position and analyzing by student’s t-test with α=0.05. ** denotes p<0.01. Each data set contains combined data for 3 biological replicates. Number of nuclei in each data set as follows: n(0kb)=922, n(1.3kb)=915, n(6.3kb)=888, n(7.6kb)=991, n(7.7kb)=681, n(8.8kb)=840, n(10kb)=809. (F) Schematic representation of distance-dependent search space model and correlation with transcriptional onset time. (G) Schematic representation of enhancer-position dependence and correlation with active state stability.

We next sought to quantify how various transcriptional parameters might be affected by each feature of E-P configuration (linear distance vs. enhancer positioning). Leveraging the dynamic temporal information encoded in the MS2 signal trajectories, we first quantified the timing of transcriptional activation across active nuclei in each E-P configuration (Figures 2B and 2C). These distributions revealed that transcriptional activation onset is delayed systematically with increased E-P distance, regardless of enhancer positioning. Thus, E-P distance governs the timing of transcription initiation independently of enhancer positioning (Figure 2C). This delay aligns with physical models where larger genomic distances expand the 3D search space required for productive E-P interactions, prolonging the stochastic collision process preceding transcriptional initiation (Figure 2F) ^28–32^. While the delay in onset time explains the moderate output reduction with distance, it fails to account for the consistently weaker activity driven by the downstream enhancer constructs relative to their upstream counterparts. This points to additional positioning-specific regulatory mechanisms beyond linear genomic spacing.

To identify parameters regulating the reduced transcriptional output in downstream enhancer configurations, we evaluated two metrics defining transcriptional competency, or the capacity to maintain stable, high-amplitude expression. “NC14 fraction active” quantifies the fraction of NC14 spent in the transcriptionally active state post initial activation, and gives a measure of expression stability, disentangling positional effects from distance-dependent initiation delays (Figures 2B and 2D). “Mean transcriptional amplitude” was used to quantify the mean signal intensity across instances of active transcriptional activity within a single nucleus over the full NC14, providing a measure of transcriptional strength during active periods (Figure 2E). Downstream enhancers significantly hindered both parameters: transcriptional signals exhibited diminished intensity (reduced amplitude) and active state persistence (lower NC14 fraction active) relative to upstream counterparts (Figures 2D and 2E). These deficiencies partitioned embryos into distinct enhancer positioning-dependent cohorts, with downstream groups showing systematically lower functional capacity. The observed positional bias implies that regulatory factors extending beyond 3D E-P contact kinetics selectively constrain downstream enhancer performance (Figure 2G). Our results highlight the multifaceted nature of the regulatory framework and reveal that E-P configurations encode distinct kinetic features to regulate transcriptional activity.

### Enhancer-promoter configuration modulates transcription independently of sequence or reporter context

Building on the finding that E-P configuration modulates transcription kinetics through two distinct modes – onset timing and active state stability – we aimed to investigate whether these regulatory effects were intrinsic to the sequences in our reporter system, or if they represented broader features of E-P configuration-dependent transcriptional control. As such, we selected a new reporter gene (intron-truncated *yellow* [*t.yel*]) and enhancer (*even-skipped* stripe 2 enhancer [eve2]) to evaluate in our system ^33^.

We generated a modified *yellow* reporter sequence (*t.yel*) by truncating 1.6 kb from its intron to match the 3 kb length of the original *lacZ* construct, eliminating gene length as a confounding factor in transcriptional signal interpretation (Figure 3A). We carried out a comparative analysis of mRNA production behavior of *t.yel* and *lacZ* reporters in the mid-range (7.6kb and 7.7kb) constructs with opposing enhancer positioning, under snaDE and evePr control. This analysis revealed that the downstream enhancer configuration still gave rise to significantly reduced mRNA production compared to the upstream counterpart (Figure 3B). Further, the activation kinetics and functional stability behaviors driven by the *t.yel* reporter gene were consistent with expected trends reported for *lacZ*. As evidenced by heat maps, regardless of reporter gene, EnhUP drove stronger, more stable transcriptional activity as compared to EnhDOWN while maintaining nearly identical t_50%_, thus supporting reporter gene neutrality in configuration-dependent effects (Figures 3C and 3D and Supplemental Figure 3). Replacing *lacZ* with *t.yel* in the 10 kb EnhDOWN construct also drove delayed onset and reduced mRNA output (Supplemental Figure 4). Notably, motif analysis revealed five CCCTC-binding factor (CTCF) binding sites (Figure5G 3A) – a core boundary element in mammalian systems – in *lacZ* compared to none in *yellow*, yet both reporters exhibited equivalent transcriptional repression in downstream enhancer configurations. This demonstrates that CTCF-mediated insulation – a proposed mechanism for positional effects – does not drive the observed enhancer positioning-dependent repression, redirecting attention to alternative regulatory mechanisms governing transcriptional competency.

**Figure 3.**
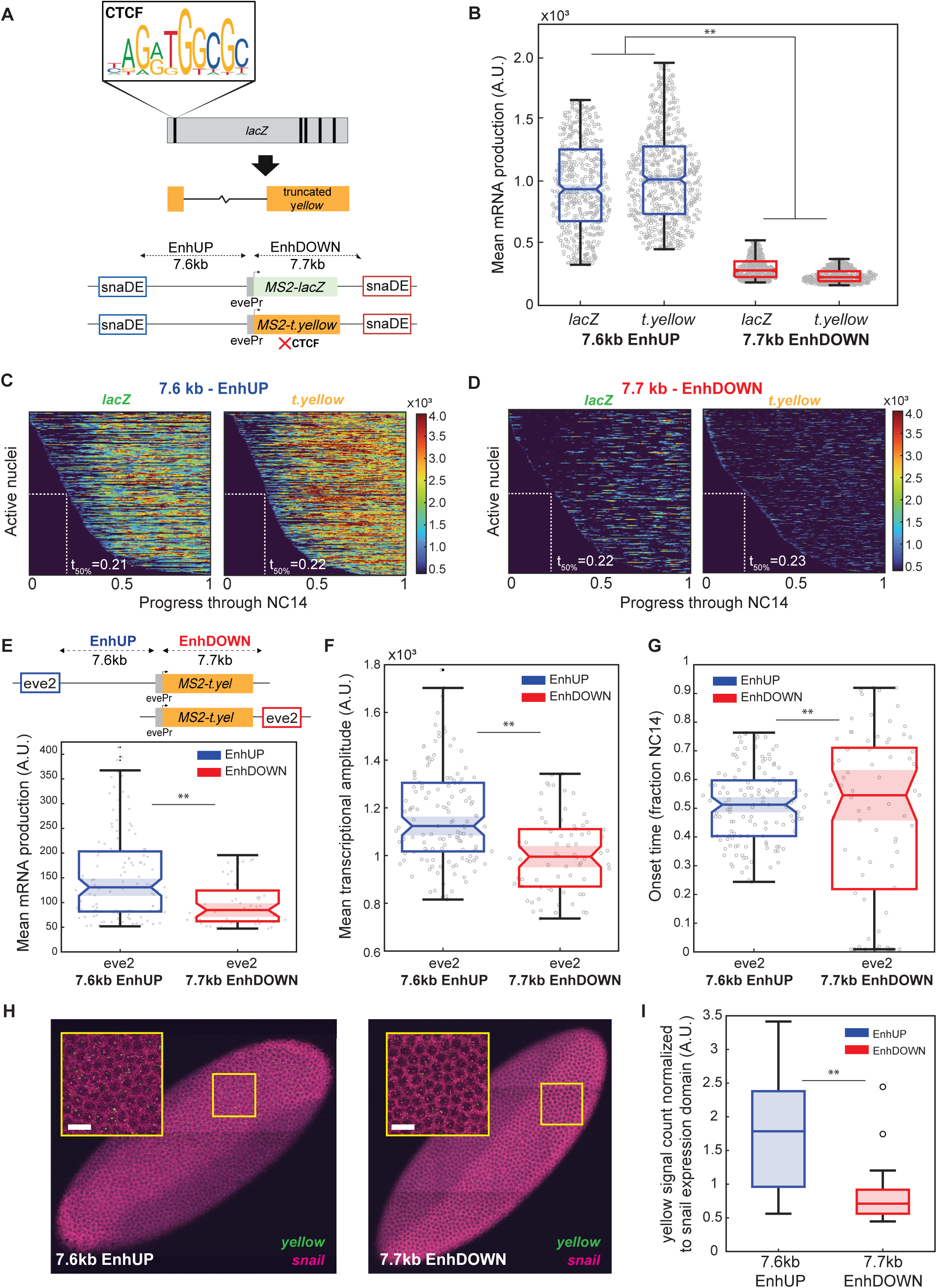
Enhancer-promoter configuration modulates transcription independently of sequence or reporter context. (A) Schematic of CTCF binding sites in *lacZ* (five) and *t.yellow* (none) and resulting construct design upon replacing *lacZ* with *t.yellow* in the 7.6 kb-EnhUP and 7.7 kb-EnhDOWN configurations. (B) Mean mRNA produced by all active nuclei in NC14 for each E-P configuration shown. Statistical significance between EnhUP and EnhDOWN subsets performed by combining the data into two groups based on position and analyzing by student’s t-test with α=0.05. Each set contains combined data for 3 biological replicates. Number of nuclei in each data set as follows: n(7.6kb-*lacZ*)=991, n(7.6kb-*t.yellow*)=909, n(7.7kb-*lacZ*)=681, n(7.7kb-*t.yellow*)=801. (C,D) Heat map of transcriptional trajectories for all active nuclei in each construct shown. The timing of activation of half the total active nuclei is reported as t_50%_. (E) Schematic representation of replacing snaDE with eve2 in the 7.6 kb-EnhUP and 7.7 kb-EnhDOWN configurations along with mean mRNA produced by all active nuclei in NC14 for these constructs. (F) Mean transcriptional amplitude of the active state plotted for each shown configuration. (G) Fraction of NC14 after onset time that each nucleus spends in the active state plotted for each shown configuration. Statistical significance for E, F, and G determined by student’s t-test with α=0.05. ** denotes p<0.01. Dots in B, E, F, and G represent total mRNA produced by individual active nuclei over the full NC14. Each set contains combined data for 3 biological replicates. Number of nuclei in each data set as follows: n(7.6kb-*eve2*)=224, n(7.7kb-*eve2*)=56. (H) RNA FISH with *yellow* probes in green and *snail* probes in magenta. Scale bar=10 μm. (I) Count of *yellow* probe signals in the entire embryo normalized to the *snail* expression domain demarcated by the high intensity cytoplasmic *snail* signal. EnhUP data contains n=13 biological replicates. EnhDOWN data contains n=15 biological replicates.

We also probed the enhancer sequence-dependence by replacing the snaDE enhancer with eve2 in mid-range *t.yel* reporter constructs (7.6 kb EnhUP and 7.7 kb EnhDOWN). While snaDE is bound by the dorsoventral (DV) regulator Dorsal, eve2 expression is controlled by anterior-posterior (AP) gap genes and drives endogenously weaker expression than snaDE ^19,34–36^. Despite these distinct genomic contexts and functional strength, the core regulatory logic persisted. Downstream configurations again showed significantly reduced mRNA production (Figure 3E), attributable to diminished transcriptional competency, as reflected by lower mean amplitudes (Figure 3F). The fold-change in mRNA output was less than that for opposing enhancer positions driven by snaDE; though this is explained by the weaker eve2 transcription strength which limits the functional range of values for all metrics. Importantly, onset delays remained the same between orientations (Figure 3G) – consistent with their near-identical linear E-P distances (7.6 vs. 7.7 kb). This replication of positioning-dependent trends across enhancers with distinct TF recruitment and endogenous genomic architectures supports that transcriptional tuning via onset timing and competency modulation is not sequence-intrinsic. Rather, these dual regulatory modes emerge as fundamental properties of E-P spatial configuration, operating independently of enhancer-specific TF binding or endogenous promoter proximity.

To validate that the observed configuration-dependent transcriptional behaviors were not artifacts of the MS2 system, where MCP-GFP binding to nascent RNA stem-loops might interfere with downstream E-P interactions, we performed independent RNA FISH experiments (Figure 3H). This fixed-tissue assay eliminates the need for MCP-GFP recruitment and directly quantifies nascent transcripts. Probes targeting the remaining intronic sequence in mid-range *t.yel* reporters with opposing snaDE positioning revealed that reduced transcriptional output in downstream enhancer configurations persisted across both imaging modalities. RNA levels, normalized to the snaDE expression domain, revealed reduced mRNA production in the downstream configuration (Figure 3I). These results support that MCP-GFP recruitment does not differentially impact transcriptional activity in upstream versus downstream positionings.

They further support that the observed transcriptional trends are not artifacts of the MS2 system, validating it as a reliable method for quantifying transcriptional dynamics at these sub-10 kb E-P configurations. Altogether, these findings highlight the robustness of both MS2-based live imaging and RNA FISH for studying transcriptional regulation and reinforce E-P configuration as a bona fide regulatory factor that influences transcription through distinct modes of tuning onset timing and active state stability.

### Two-state model of transcription reveals relative enhancer positioning is a regulator of bursting frequency

Transcriptional activity exhibits stochastic dynamics characterized by intermittent active and inactive periods, referred to as transcriptional bursting ^37,38^. We employed a two-state stochastic model of transcription to quantify bursting parameters modulated by opposing enhancer configurations (Figures 4A and 4B). Widely used to characterize transcriptional bursting, two-state models assume that the promoter state switches between the ON and OFF state with certain transition kinetics. The transition rate can be estimated based on individual transcriptional trajectories, providing estimates of two key parameters of transcriptional bursting: burst amplitude and frequency ^39–42^. Burst amplitude refers to the average number of mRNA molecules produced during a transcriptionally active period, and burst frequency represents the rate at which the gene switches from an inactive to active transcriptional state. These parameters exhibit variability across genomic contexts and are influenced by the regulatory element composition of individual transcriptional loci ^43^. We investigated whether three-dimensional configurational parameters – operating within identical regulatory sequences – exert additional control over transcriptional bursting dynamics.

**Figure 4.**
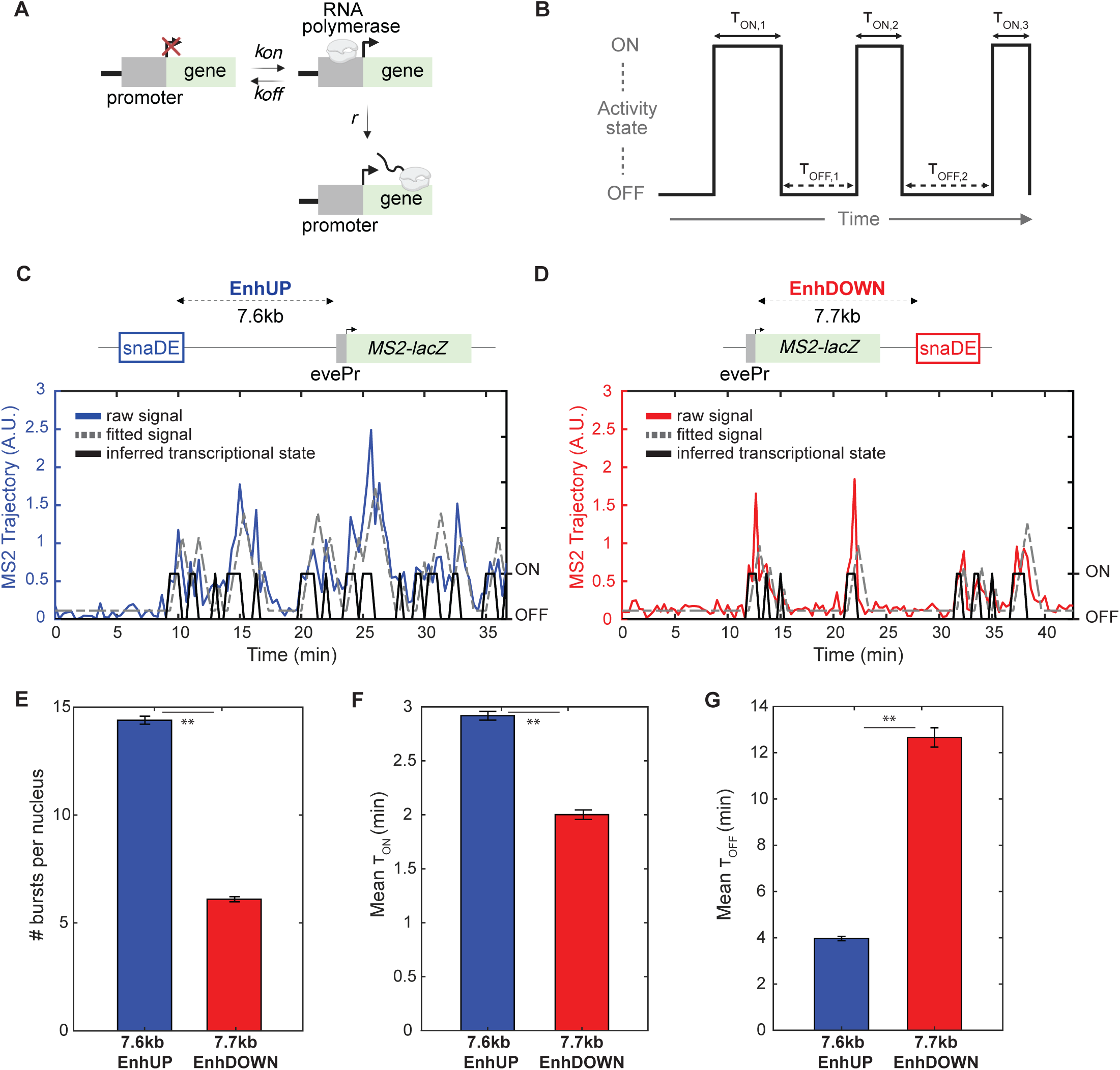
Two-state model of transcription reveals relative enhancer positioning is a regulator of bursting frequency. (A) Schematic of promoter activity model. (B) Representative trajectory of inferred promoter state where ON and OFF time periods are defined as τ_ON_ and τ_OFF_, respectively. (C, D) Raw trajectory, predicted trajectory, and predicted activity state plotted over full NC14 for EnhUP (C) and EnhDOWN (D) configurations. (E) Number of bursts per active nucleus. (F) Mean τ_ON_ for all bursts in each active nucleus. (G) Mean τ_OFF_ between all bursts in each active nucleus. ** denotes p<0.01 for student’s t-test conducted with α=0.05. Error bars represent standard error of the mean. Each set contains combined data for 3 biological replicates. Number of nuclei in each data set as follows: n(7.6kb) =991 and n(7.7kb)=681.

The model was applied to mid-range constructs featuring opposing enhancer orientations, 7.6 kb-EnhUP and 7.7 kb-EnhDOWN, respectively (see Methods for modeling details) ^44^. Our model accurately recapitulated the stochastic transcriptional dynamics from both constructs, showing strong agreement with experimental MS2 fluorescence intensity trajectories (Figures 4C and 4D). From the transcriptional trajectories of individual nuclei, we measured the duration of each burst (τ_ON_) and the time interval between successive bursts (τ_OFF_) (Figure 4B). The number of bursts across individual trajectories was reduced more than 50% in the EnhDOWN compared to EnhUP constructs – consistent with the reduced transcriptional stability reported previously (Figure 4E). Furthermore, the duration of each burst (denoted as τ_ON_) was shortened in the EnhDOWN configuration, contributing to the lower burst amplitude for EnhDOWN configurations reported in earlier sections (Figure 4F). A more significant difference was found in the time interval between successive bursts (τ_OFF_), where the EnhDOWN configuration induced an approximately 3-fold increase compared to EnhUP (Figure 4G). This supports the hypothesis that downstream enhancer positioning destabilizes transcriptionally active states, contributing to a shorter burst duration and longer inter-burst periods. Smaller burst size and reduced burst frequency resulted in the 70% decrease in mRNA production observed in the EnhDOWN construct compared to the EnhUP construct (Figure 1F). Crucially, this divergent kinetic behavior emerged from identical snaDE sequences and reporter components differing only in their relative genomic positioning (upstream vs. downstream). These findings establish that relative enhancer positioning at sub-10 kb scales directly alters transcriptional bursting kinetics, implicating E-P configuration – independent of sequence variation – as a deterministic regulator of mRNA production dynamics.

### Locus-associated Dorsal hub behaviors are sequence-intrinsic and independent of enhancer-promoter configuration

To further elucidate the mechanistic basis of E-P configuration-dependent transcriptional regulation, we analyzed if opposing enhancer configurations might engage distinct *trans*-regulatory elements – such as transcription factor hubs – that provide feedback to stabilize transcriptional states.

The TF hub model proposes that dynamic, spatially restricted clusters of TFs, coactivators, and RNA polymerase II form near active promoters and function as self-renewing reservoirs of transcriptional machinery ^45–47^. Leading models suggest these hubs stabilize productive E-P interactions through multivalent interactions involving intrinsically disordered regions ^48,49^. While debates persist about the regulators of hub formation and composition, mounting evidence indicates that *cis*-regulatory element sequences – particularly enhancer grammar and density – direct differential hub formation ^50,51^. Given that our experimental system reveals striking transcriptional stability differences based on enhancer positioning (upstream vs. downstream), a key mechanistic question arises: do configurational parameters like relative enhancer position influence hub dynamics or are hub behaviors instead governed by sequence-intrinsic properties of enhancers, independent of genomic positioning? Resolving this dichotomy might clarify how spatial genome organization interfaces with local transcriptional clusters to regulate gene expression.

If E-P configuration modulates transcription factor hub behavior, we predict distinct hub dynamics at transcriptionally active loci in EnhUP versus EnhDOWN reporter configurations. To test this, we implemented a dual live-imaging platform combining endogenously tagged Dl-GFP (a core TF binding the snaDE enhancer) with the MS2/MCP-mCherry system for real-time transcription visualization under a high-resolution lattice light-sheet microscope (Figures 5A and 5B). This enabled simultaneous 3D tracking of both Dl clusters – serving as a functional proxy for whole-hub behavior – and nascent *lacZ* reporter transcripts in mid-range E-P configurations (7.6kb-EnhUP and 7.7kb-EnhDOWN) regulated by snaDE, thus enabling quantification of hub behaviors and any correlations with transcriptional activity or enhancer-promoter configuration.

**Figure 5.**
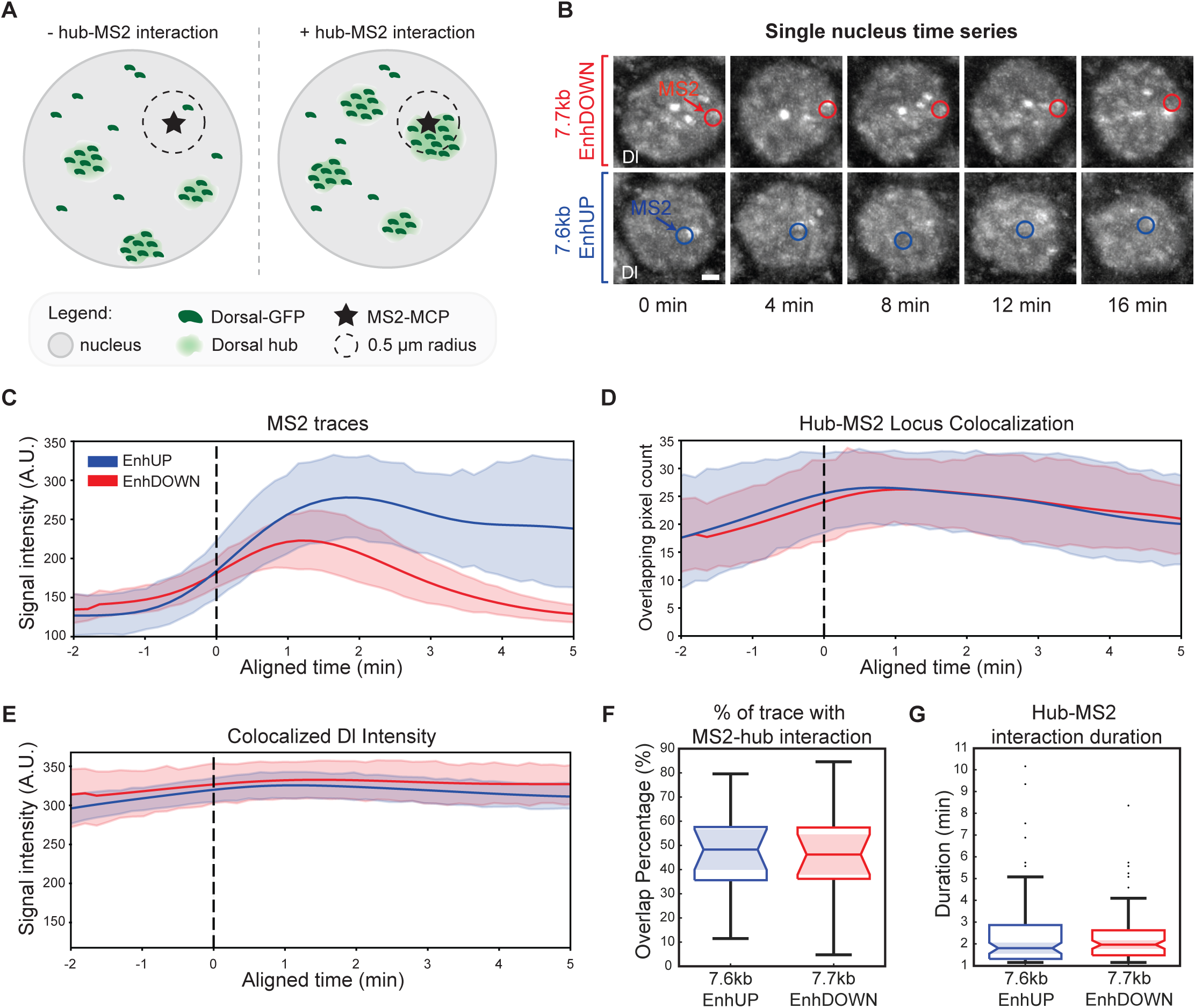
Locus-associated Dorsal hub behaviors are sequence-intrinsic and independent of enhancer-promoter configuration. (A) Schematic representation of non-MS2 associated Dorsal hubs (left) compared to locus-associated Dorsal hubs within the 0.5 μm radius sphere centered on the transcription locus. (B) Time series snapshots of an active nucleus over time for EnhDOWN (top) and EnhUP (bottom) constructs. The transcription locus is circled and labeled <MS2>. Dorsal protein is visualized in gray. Scale bar=1 μm. (C) Mean smoothed transcriptional trajectory for all active nuclei. (D) Count of Dl hub voxels overlapping with 0.5 μm radius sphere centered on the transcription locus. (E) Mean intensity of Dl hub pixels within 0.5 μm radius sphere centered on the transcription locus. Plots C-E are aligned on the x-axis to each nucleus’s transcription onset time where onset time is set to time zero on the aligned plot. (F) Percentage of each nucleus’s MS2 trace during which a proximal Dl hub shared pixels with the 0.5 μm radius sphere centered on the transcription locus. (G) Duration in minutes of each Dl hub-MS2 colocalization event. Student’s t-test with α=0.05 for F and G reported no statistically significant differences between EnhUP and EnhDOWN in either panel. Data in panels C-G analyzed from n=16 nuclei for each construct.

TF hub engagement was quantified by analyzing the Dl-GFP clusters within a 0.5 μm radius of active transcription sites. Loci meeting this spatial criterion were operationally classified as “transcriptionally engaged” (Figure 5A). We quantified two hub behavior metrics: the mean Dl concentration of hubs engaged with active *lacZ* transcription loci (Colocalized Dl Intensity) and the number of hub voxels colocalizing with the 0.5 μm transcription zone (Hub-MS2 Locus Colocalization). Temporal analysis throughout NC14 revealed no significant differences in either parameter between EnhUP and EnhDOWN configurations (Figures 5D and 5E), despite their divergent transcriptional output evidenced by MS2 intensity traces (Figure 5C). To address whether enhancer positioning modulates TF hub interaction kinetics, we further analyzed the probability of Dl hub-MS2 contact by quantifying the percentage of NC14 that each active locus spent in a locus-associated hub interaction (% of trace with MS2-hub interaction) (Figure 5F). Additionally, the persistence of hub-MS2 contact was quantified by measuring the duration of contact between MS2 and locus-associated hubs (Hub-MS2 interaction duration) (Figure 5G). Similar to the other metrics, we observed no significant difference in Hub-MS2 interaction persistence or probability across constructs.

These results demonstrate a clear dissociation between TF hub dynamics and E-P configuration-dependent transcriptional activity. Transcription factor hub formation, spatial organization, and functional engagement appear to be principally governed by *cis*-regulatory sequence features (enhancer/promoter identity), with no detectable influence from E-P configuration mechanisms in this system.

## Discussion

In this study, we uncover a previously uncharacterized layer of *cis*-regulatory control in which two parameters of E-P configuration – linear distance and relative positioning – govern distinct aspects of transcriptional kinetics at the sub-10 kb genomic length scale. Through systematic designs, quantitative single-cell live imaging, and mathematical modeling, we demonstrate two distinct regulatory effects: linear E-P distance determines the timing of transcriptional activation, while enhancer positioning (upstream vs. downstream) dictates the stability of the transcriptionally active state post-initiation, resulting in divergent transcriptional bursting dynamics and mRNA output. Thus, our findings challenge conventional models of enhancer logic where enhancers function independently of relative positioning, by highlighting local E-P configuration as a key driver of transcriptional regulation at short genomic distances.

The observed distance-dependence on transcriptional output likely arises from a diffusion-controlled model wherein increased spatial separation in the post-mitotic, relaxed chromatin state expands the three-dimensional search space required for cognate regulatory elements to establish productive interactions. We propose that this mechanism may play a role in regulating proper spatiotemporal gene expression, for example, by ensuring specific, sequential reactivation of Hox genes regulated by a single shared enhancer after mitosis, thus maintaining positional identity in rapidly dividing progenitor cells ^52^.

Further, we uncovered a surprising result that relative E-P positioning affects gene expression by regulating transcriptional stability. This raises an important question: does the inherent stability challenge associated with downstream enhancer positioning serve a specific regulatory function by promoting reduced, more variable transcription? Or is it simply a biophysical consequence that endogenously downstream-positioned enhancers must overcome through compensatory mechanisms (such as recruiting tethering elements or *trans*-acting factors)? Our findings suggest that this positioning bias could serve as an inherent timing mechanism during early embryonic development, where transient transcriptional stability windows must align with developmental milestones. For instance, in *Drosophila* embryogenesis, the positioning of pair-rule gene enhancers relative to their promoters may encode burst duration thresholds that coordinate segmental patterning ^39,53^. The prevalence of downstream enhancers in endogenous contexts makes this line of inquiry particularly critical for understanding transcription regulation *in vivo* and merits further investigation to more thoroughly characterize the functional importance of E-P configuration-dependent transcription regulation.

In probing the underlying mechanism of enhancer position-dependent transcriptional regulation, we studied the dynamics of locus-associated Dorsal hubs under opposing enhancer positions. Our analysis of hub interactions with active transcriptional loci revealed no direct correlation between transcriptional activity and hub-loci association dynamics; though our analysis was limited only to active transcriptional windows, thus we cannot discount that TF hub formation probability may be affected by E-P configuration parameters. This supports the model that TF hub formation is a sequence-intrinsic property, consistent with recent studies ^51^. However, we note that tracking a single TF (Dorsal) as a proxy for hub behavior may not be representative of the diverse composition of the TF hubs in this system. Consequently, our findings do not fully rule out transcription-dependent modulation of hubs mediated by other low-abundance or weakly binding factors, which could couple mRNA production to transcriptional state stability. Further studies will be necessary to explore the functional interplay, if any, between TF hubs and E-P configuration parameters by visualizing other TFs or Pol II.

While simultaneous visualization of hubs and transcription introduces technical constraints, our data suggest that TF hubs are unlikely to underlie our observation of positioning-dependent transcriptional stability. Instead, we propose a mechanical model in which DNA strand tension across sub-10 kb E-P distances constrains chromatin’s capacity to sustain looped configurations required for stable transcription. In downstream enhancer (EnhDOWN) configurations, transcription-generated mechanical stress – driven by DNA uncoiling and RNA polymerase II (Pol II) procession – amplifies strand tension ^54–56^. This elevates the thermodynamic barrier to maintaining E-P proximity, favoring rapid interaction dissociation and transcriptional termination. This model reconciles the observed positioning-independent, post-mitotic activation timing, which depends solely on the linear E-P distance. Before transcription activation, baseline tension is comparable in both EnhUP and EnhDOWN configurations, allowing linear E-P distance to dictate activation kinetics. Post-initiation, however, position-specific tension differentially affects EnhUP vs. EnhDOWN E-P interactions, leading to divergent transcriptional stability. Future studies will directly test this hypothesis by modulating strand tension (e.g., by varying the effective reporter gene length) within the sub-10 kb E-P regime, evaluating its role as a mechanochemical regulator of short-distance transcriptional dynamics.

This work reveals an additional, nuanced layer of regulatory logic whereby transcriptional regulation is governed by two sequential, spatially constrained processes: enhancers make initial contact with the target promoter and the interactions are then stabilized to facilitate transcription in an E-P configuration dependent manner. While these mechanisms dominate at short length scales, their relative contribution to gene regulation remains undefined in the endogenous context where long-range chromatin architecture regulation is also at play. To resolve this regulatory hierarchy, future studies should systematically dissect how active chromatin remodeling systems interface with local E-P configuration parameters. Incorporating insulator or tethering elements into our modular experimental platform would enable quantitative mapping of how these chromatin remodelers compete with or reinforce configuration-dependent E-P interaction kinetics and define their role in fine-tuning transcriptional output at sub-TAD scales.

The integration of spatial E-P configuration rules with mechanochemical feedback loops – which convert chemical input (e.g., TF and Pol II recruitment to promoter region) into mechanical feedback (e.g., increased strand tension influencing E-P interactions) – provides a robust framework for understanding developmental precision in gene regulation. By constraining transcriptional output through biophysical parameters rather than (or as well as) sequence-specific codes of enhancer syntax, evolution could tinker with enhancer placement to fine-tune developmental timing without compromising core regulatory logic, thus ensuring robust and reproducible phenotypes. This paradigm shift has far-reaching implications for understanding complex genetic disorders and developing targeted therapeutic strategies, potentially revolutionizing our approach to precision medicine and genetic engineering.

## Resource Availability

Available upon request.

## METHODS

### Fly strains generated via PhiC31 Integration

In all experiments, *Drosophila melanogaster* embryos were studied during the entire NC14 from mitosis to gastrulation. All plasmids containing a reporter gene of interest were integrated to the *vk33* landing site on the third chromosome (Bloomington *Drosophila* Stock Center, cat# 9750) by micro-injection performed at BestGene, Inc. Virgin progeny from the nos>MCP-GFP, His-RFP and yw cross were crossed with males carrying the MS2 reporter genes^18^. Embryos from the cross were imaged under a confocal microscope for MS2-only studies and a lattice light-sheet microscope for the MS2 and Dorsal simultaneous visualization studies.

### Plasmid construction

All plasmids were constructed by insertion of enhancer, promoter, reporter, and MS2 sequences to a backbone containing the *miniwhite* marker, an attB recombination site, and ampicillin resistance gene.

#### Enhancer and promoter sequences

For all constructs, we used the 126 bp minimal *even-skipped* promoter (evePr) to drive transcription of the MS2-*lacZ* reporter gene. Two different enhancer sequences were analyzed in this study. These are the 1.5 kb *snail* distal enhancer (snaDE) and the 728 bp *even-skipped* stripe 2 enhancer (eve2).

#### Reporter genes

In this study, we utilized the 3 kb *lacZ* gene and the 3 kb intron-truncated *yellow* gene as reporters. The intron-truncated length normalization prevented artificial inflation of MCP-GFP signals caused by prolonged Pol II residence time on longer templates, which could otherwise mimic elevated mRNA production through extended MS2 RNA stem-loop visibility. Additionally, the L-tubulin 3’UTR was located downstream of each reporter gene in all constructs.

#### Neutral spacer sequences

In this study, two neutral spacer sequences were utilized to vary the linear enhancer-promoter distance in the reporter constructs. One sequence, *yeeJ-1*, is a 5 kb fragment of the E*. coli* gene *yeeJ.* Site directed mutagenesis was performed to remove a Zelda binding site from the sequence. PCR primers for *yeeJ-1* amplification and Zld SDM are provided below. A second neutral spacer sequence was computationally designed to lack TFs relevant in early *Drosophila* development ^21^.

#### Screening for TF binding motifs in spacer and reporter sequences

In this study, we use consensus binding motifs of key transcription factors to evaluate the likelihood of putative TF binding sites in our neutral spacers and reporter sequences. The binding motifs were obtained from the JASPAR database and downloaded in MEME format. The FIMO program from MEME-suite version 5.5.7 was used to identify high-probability (p<0.0001) binding sites for the screened TFs (Dl, Zld, Twi) in each of the spacer and reporter sequences. A summary of the results can be found below in Table 1.

**Table 1.**
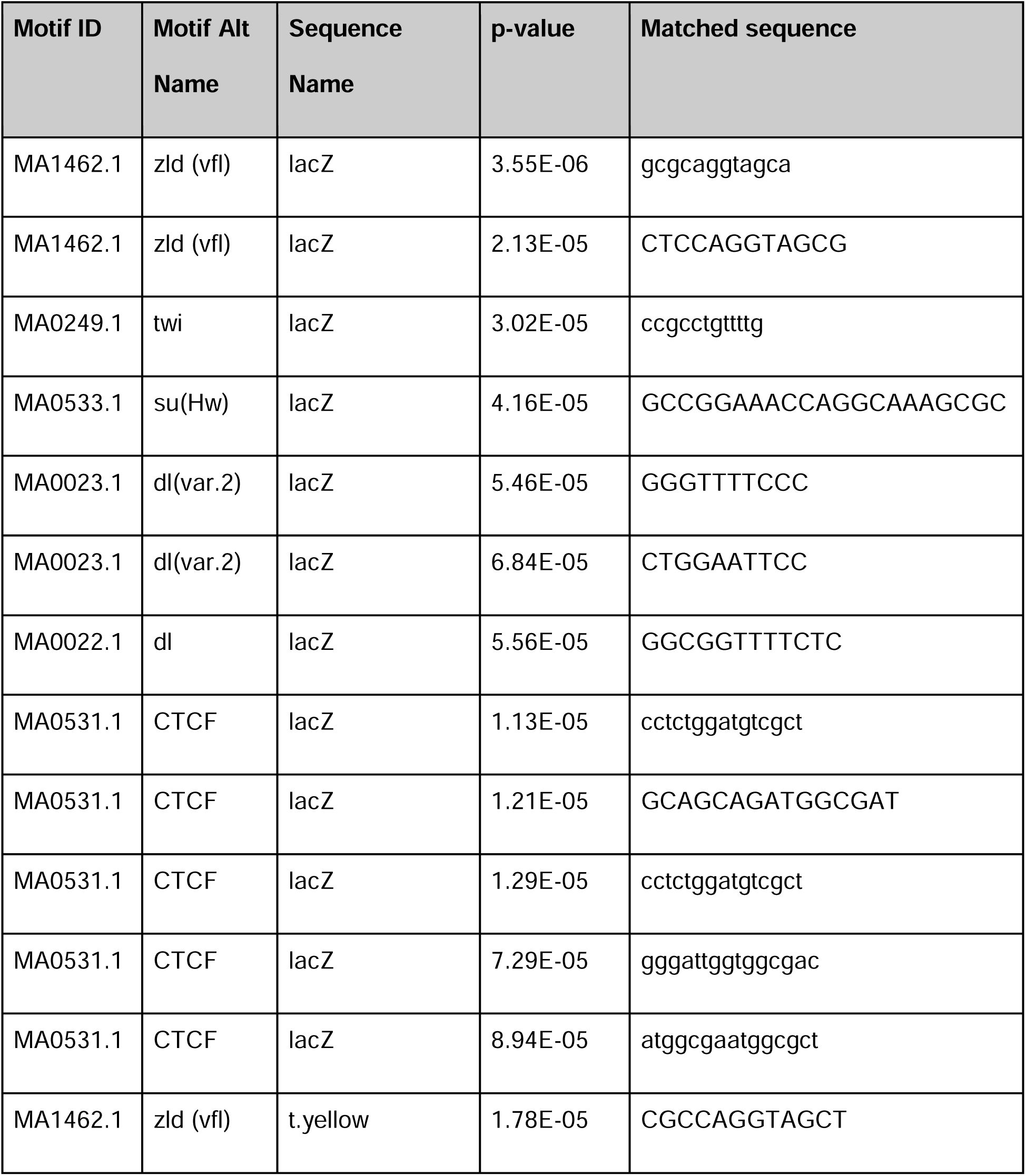
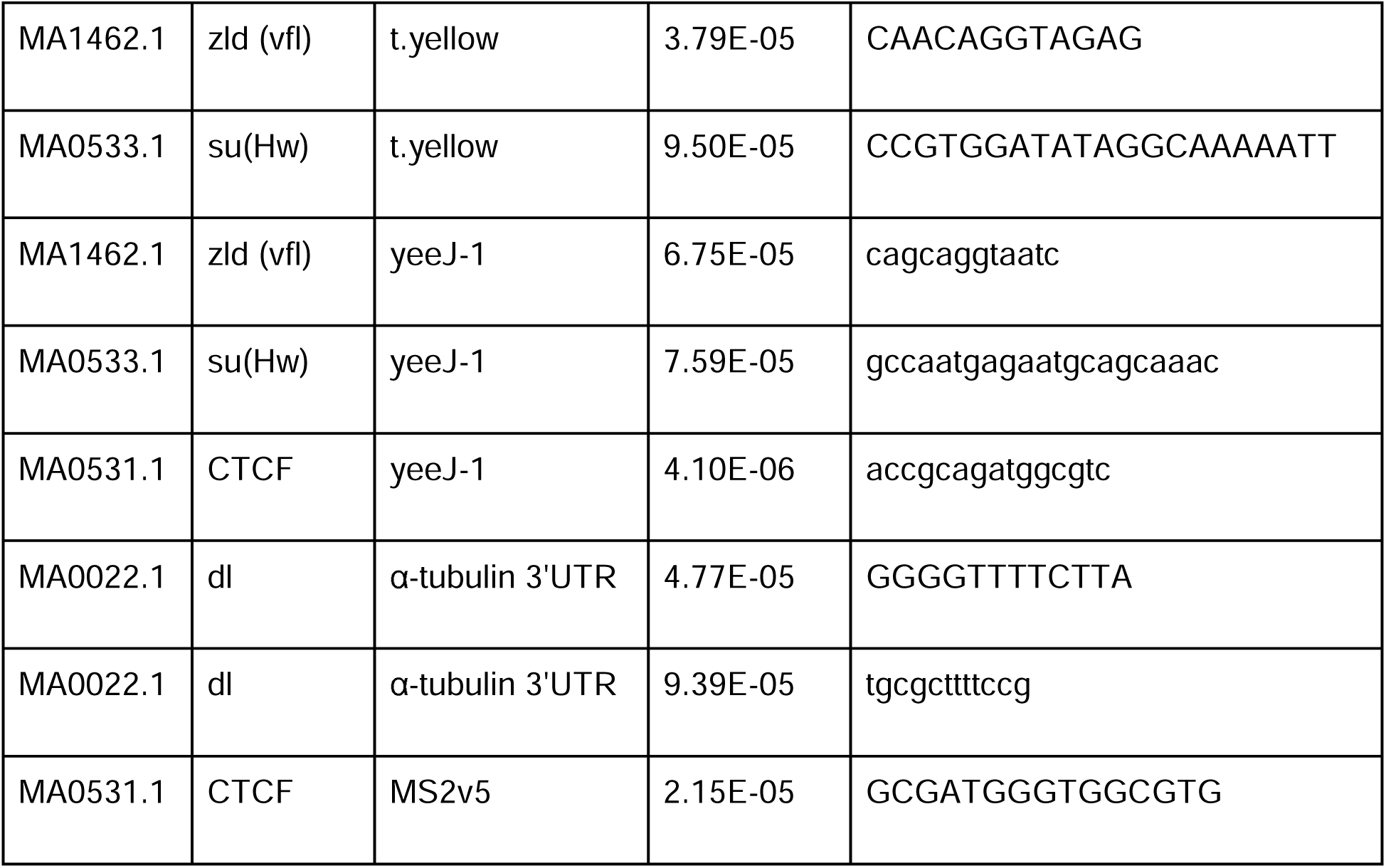
Binding motif identification in neutral spacer and reporter sequences.

### Generation of endogenous Dorsal-eGFP fly line

Dorsal-eGFP was generated using CRISPR-cas9 homology-directed repair to insert a poly-glycine/serine linker+eGFP sequence into the 3’ end of the endogenous dorsal locus. Two target sites within the fifth dorsal intron and 3’UTR were selected using the Target Finder (http://targetfinder.flycrispr.neuro.brown.edu) tool.

The annealed Dorsal gRNA oligos were subcloned into the pCFD3-dU6:3gRNA vector (a gift from Simon Bullock, Addgene plasmid #49410; http://n2t.net/addgene:49410; RRID:Addgene_49410) via BbsI restriction sites. Approximately 1-kb fragments of dorsal homology arm sequences were synthesized and inserted into the CRISPR-eGFP-loxP-dsRed-loxP plasmid. To create the linker+eGFP 3’ fusion the dorsal genomic region in between the guide sequences was obtained via PCR and inserted with the linker+eGFP into the CRISPR-eGFP-loxP-dsRed-loxP plasmid via Gibson Assembly. The dorsal stop codon was replaced with a stop codon at the 3’ end of eGFP sequence for proper translation. The gRNAs and dorsal homology arm eGFP plasmids were co-injected into nos-Cas9 embryos (TH00787.N), and DsRed+ progeny were screened (BestGene, Inc.). The resulting endogenous dorsal-eGFP fly strains were homozygous viable.

## Quantification And Statistical Analysis

### Live imaging MS2 activity only

Virgin progeny from the nos>MCP-GFP, His-RFP and yw cross were crossed with males carrying the MS2 reporter genes ^18^. Embryos from the cross were dechorionated and mounted in Halocarbon oil (Sigma) between a semipermeable membrane (Sarstedt) and coverslip (18 mm x 18 mm). Embryos were imaged using a Zeiss LSM800 and Plan-Apochromat 40x/1.3 numerical aperture (N.A.) oil-immersion objective. A single image was 512 x 512 pixels. Images were created by taking a maximum projection from a stack of 14 images separated by 0.75 μm, and the final time resolution was 20 s per frame. Image capture was started from the beginning of NC14 mitosis to gastrulation. Images were captured in 16-bit. Since the eve2-driven constructs exhibited too low transcriptional activity to be captured by the same laser power as the snaDE-driven constructs, we used two different laser powers that were optimized for each construct. Importantly, the same laser power was used for all snaDE constructs and a different, consistent laser power was used for all eve2 constructs.

### Image analysis for MS2 activity only

Segmentation and tracking of nuclei were carried out following the methodology outlined in (REF).. Every Drosophila strain used in this research expressed His2Av-eBFP2 to label nuclei. For live image processing and nuclei segmentation, a Gaussian filter was initially applied to the nuclei-labeled channels to smooth out the images. This was followed by top-hat filtering and contrast-limited adaptive histogram equalization to correct for uneven illumination and to improve image contrast. The processed images were then binarized using a threshold determined by Otsu’s method. Manual corrections were made to the binary images of each frame to ensure segmentation accuracy. Each nucleus identified in the segmentation was assigned an index, and the x and y coordinates of all constituent pixels, as well as the center of mass, were documented. A custom MATLAB script was used to track segmented nuclei from nuclear cycle 14 up to the beginning of gastrulation. To track each indexed nucleus, the Euclidean distance between a nucleus and all nuclei in the following frame was computed and compared. Nucleus pairs with the smallest Euclidean distance, provided it was less than 10 pixels between consecutive frames, were matched and considered to represent the same nucleus over time. If the minimum distance exceeded 10 pixels, the nucleus was classified as having no corresponding lineage, due to partial or complete movement out of the imaging frame. For snaDE-driven constructs, MS2 signal intensities were extracted by taking the average of the highest two pixels from each nucleus using a mask created by segmenting the embryo. For eve2-driven constructs, due to weaker intensities and reduced signal:noise ratio, a modified MS2 extraction algorithm was used. The 9-pixel average intensity centered around the 2 brightest pixels within each nucleus mask was calculated and the highest of the two was selected as the region with “true” signal. Of those 9-pixels in the “true” signal region, the average of the brightest 2 was used to represent the MS2 signal intensity. Background subtraction for the MS2 channel was performed by subtracting the channel’s mean intensity of initial non-transcribing signals within all nuclei of an individual embryo. The traces were then normalized by dividing each point by the maximum value within each trace. For snaDE-driven constructs, the mRNA output was measured by taking the area under the curve of each MS2 trajectory over the entire NC14 time course. For eve2-driven constructs, again due to weak signal and poor signal:noise ratio, the mRNA output was measured by integrating under the curve only during frames of active transcription (i.e. signal intensity above an empirically defined threshold value corresponding to visible MS2 puncta).

#### Live imaging Dl hub and MS2 simultaneously

Virgin progeny from the Dl-eGFP and nos>MCP-mCherry cross were crossed with males carrying the MS2 reporter genes. Embryos were collected over a 90 minute window and manually dechorionated using double-sided tape and a blunt needle. Dechorionated embryos were mounted on agar pads and transferred to a heptane-acrylic adhesive on 25 mm glass coverslips, created by dissolving double-sided tape in heptane.

Imaging was performed using a custom-built lattice light-sheet microscope^57^, based on an adaptive optics configuration originally developed by the Betzig lab^58^. Excitation was achieved using 488 nm and 589 nm laser lines. Beams were expanded to 2 mm, polarized via a half-wave plate, and directed into an acousto-optic tunable filter, or AOTF (Quanta-Tech, AA Opto Electronic), for wavelength selection and power modulation. A Powell lens (Laserline Optics Canada) and cylindrical lenses shaped the beam, which was relayed through a second half-wave plate to a spatial light modulator (Meadowlark Optics, AVR Optics) and projected onto a custom annular mask. The resulting beam was relayed through a resonant galvanometer and a pair of galvo scanning mirrors (Cambridge Technology, Novanta Photonics) before being focused onto the sample with a Thorlabs TL20X-MPL objective. Fluorescence was collected orthogonally by a Zeiss 20× 1.0 NA objective and relayed to a deformable mirror (ALPAO) for adaptive correction. Emission was split by a dichroic (Semrock Di03-R561-t3-25x36) and imaged onto two sCMOS detectors (Hamamatsu ORCA Fusion) with emission and notch filters appropriate for GFP (Semrock FF03-525/50-25 emission filter and a Chroma ZET488NF) and mCherry (Semrock FF01-615/24 emission filter and a Chroma ZET594NF notch filter). A multi-Bessel lattice sheet (NA 0.4/0.3) was used for excitation. Volumes spanning 18.9 µm were captured at 0.3 µm z-steps with 30 msec exposures per plane and 9.31-second intervals between stacks. Laser power was 0.124 mW for 488 nm and 1.344 mW for 589 nm.

### Image analysis for Dl hub and MS2 simultaneous visualization

All captured images were deconvolved using GPU-accelerated Richardson-Lucy deconvolution with 8 iterations and bead-derived point spread functions^59^. For visualization, TIFF and Zarr images were converted to Imaris format using a custom converter^59^. Nuclear segmentation was performed using a custom-trained Cellpose 2.0^60^ model, trained on MCP-mCherry-labeled embryos using manually curated micro-SAM^61^ data. Segmented slices were reconstructed into 3D masks with post-processing steps for stitching and gap interpolation. Nuclear tracking across time was performed using a nearest-neighbor approach. Hub segmentation was performed using a pipeline previously described^51^. Nuclei were normalized to their mean intensity, and hub regions were identified using median filtering, morphological reconstruction, and subtraction.

Local maxima were used as seeds for watershed segmentation, enabling resolution of closely packed or amorphous hubs. Region properties including integrated intensity, size, and mean intensity were extracted.

MS2-positive nuclei were manually selected based on segmentation quality and MS2 spot visibility. For each selected nucleus, a 4D crop was generated. MS2 spots were segmented using a combination of median filtering, difference-of-Gaussians, and adaptive thresholding. Parameters were tuned per nucleus to optimize segmentation. The brightest spot within each nucleus was tracked over time using nearest-neighbor tracking with manual corrections. MS2 positions were extrapolated 20 frames before burst onset and interpolated across gaps due to transcriptional pulsing. To quantify hub-locus interactions, we calculated the overlap between segmented hubs and a 0.5 µm sphere centered at the MS2 centroid as performed previously^50^. Interaction duration, intensity, and volume were recorded.

### Fluorescence *in situ* Hybridization (FISH)

Embryos were collected 3h after laying and fixed with formaldehyde. RNA FISH followed established protocols (Lim et al., 2015) using probes specific to *sna* (BIOT & DIG) and *yellow* (FITC). The primary antibodies sheep anti-DIG, rabbit anti-FITC, and mouse anti-Biotin were applied, followed by Alexa Fluor-conjugated donkey secondary antibodies (488 anti-rabbit, 555 anti-sheep, 647 anti-mouse). Nuclei were stained with DAPI. The yellow probe was used for snaPr>MS2-yellow detection. Embryos were imaged using a Zeiss LSM800 confocal microscope (Plan-Apochromat 20×/0.8 NA) at 512×512 pixel resolution. Analyses included only ventral-oriented embryos at mid-nuclear cycle 14. Sample sizes were 13 enh-UP and 14 enh-DOWN embryos.

Mathematical model of fluorescence dynamics and transcription state inference:

Given the possible transcriptional states (ON and OFF), we define the function *ϕ_c_* as follows:

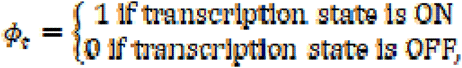

The experimental signal *y_t_* is fitted to a signal 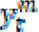 that follows our model. In our model, signal increases and decreases following linear function of time with a fitted rate λ following the difference equation in discrete time:

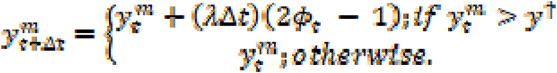

Where 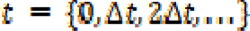 with Δ*t* = 20 *s*. The burst threshold *y*† is defined such that below this threshold, we assume that no bursting occurs and the transcription state is OFF. Our algorithm optimizes the accumulation/decay rate is optimized by the minimization of the distance between the actual fluorescence trajectory and the modeled one.

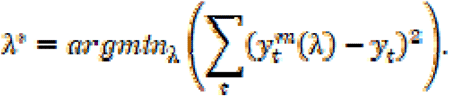

The optimal *λ^s^* will predict its respective *ϕ_t_* which is taken as the most accurate trajectory of the transcriptional state. The bursting properties are estimated based on the duration of the states on and off for the inferred *ϕ_t_*.

Statistical significance:

Sample sizes are indicated in the figure legends along with statistical significance parameters for each data set. For comparison between two independent samples, a two-tailed Student’s t-test was used with a significance threshold of α = 0.05 and significance was defined as p < 0.01. For multi-group comparisons, significance was tested by the analysis of variance (ANOVA) with a significance threshold of α = 0.05 and significance was defined as p < 0.01. All statistical analyses were conducted using MATLAB (version R2021b, Mathworks).

## Acknowledgements

We thank Lim lab members, Mike Levine, and Mounia Lagha for helpful discussions and feedback on the manuscript. We also thank Arush Arora for his help in automating the image analysis pipeline. This work was supported by National Institutes of Health grant R35GM133425 (B.L.) and DP2HD108775 (M.M.), Margaret Q Landenberger Foundation (M.M.), Howard Hughes Medical Institute, Freeman Hrabowski Scholars Program (M.M)., NIH-NIGMS grant R35GM148351(A.S.), National Science Foundation Graduate Research Fellowship grant DGE-2236662 (to S.F.), and the University of Pennsylvania Fontaine Society (E.A.L.P).

## Author contributions

E.A.L.P., S.F., M.M, and B.L. designed the experiments. E.A.L.P wrote the paper. E.L.P, J.W., S.F., E.D., R.T.T., J.A.V. conducted experiments. C.N. and A.S. performed stochastic modeling.

## Supplementary information

Document S1. Figures S1–S4

## References

1. Alexander, J. M. et al. Live-cell imaging reveals enhancer-dependent Sox2 transcription in the absence of enhancer proximity. Elife 8, e41769 (2019).

2. Wang, Z., Luo, S., Zhang, Z., Zhou, T. & Zhang, J. 4D nucleome equation predicts gene expression controlled by long-range enhancer-promoter interaction. PLoS Comput Biol 19, e1011722 (2023).

3. Zuin, J. et al. Nonlinear control of transcription through enhancer–promoter interactions. Nature 604, 571–577 (2022).

4. Banerji, J., Rusconi, S. & Schaffner, W. Expression of a β-globin gene is enhanced by remote SV40 DNA sequences. Cell 27, 299–308 (1981).

5. Gillies, S. D., Morrison, S. L., Oi, V. T. & Tonegawa, S. A tissue-specific transcription enhancer element is located in the major intron of a rearranged immunoglobulin heavy chain gene. Cell 33, 717–728 (1983).

6. Pennacchio, L. A., Bickmore, W., Dean, A., Nobrega, M. A. & Bejerano, G. Enhancers: five essential questions. Nat Rev Genet 14, 288–295 (2013).

7. Schirm, S., Jiricny, J. & Schaffner, W. The SV40 enhancer can be dissected into multiple segments, each with a different cell type specificity. Genes Dev. 1, 65–74 (1987).

8. Weber, F. & Schaffner, W. Simian virus 40 enhancer increases RNA polymerase density within the linked gene. Nature 315, 75–77 (1985).

9. Kvon, E. Z. et al. Genome-scale functional characterization of Drosophila developmental enhancers in vivo. Nature 512, 91–95 (2014).

10. Fujioka, M., Emi-Sarker, Y., Yusibova, G. L., Goto, T. & Jaynes, J. B. Analysis of an even-skipped rescue transgene reveals both composite and discrete neuronal and early blastoderm enhancers, and multi-stripe positioning by gap gene repressor gradients. Development 126, 2527–2538 (1999).

11. Han, J., Zhang, Z. & Wang, K. 3C and 3C-based techniques: the powerful tools for spatial genome organization deciphering. Molecular Cytogenetics 11, 21 (2018).

12. Raj, A. Single-Molecule RNA FISH. in Encyclopedia of Biophysics (ed. Roberts, G. C. K.) 2340–2343 (Springer, Berlin, Heidelberg, 2013). doi:10.1007/978-3-642-16712-6_518.

13. Raj, A., van den Bogaard, P., Rifkin, S. A., van Oudenaarden, A. & Tyagi, S. Imaging individual mRNA molecules using multiple singly labeled probes. Nat Methods 5, 877–879 (2008).

14. Georgiev, P. & Maksimenko, O. Mechanisms and proteins involved in long-distance interactions. Front. Genet. 5, (2014).

15. Pachano, T., Haro, E. & Rada-Iglesias, A. Enhancer-gene specificity in development and disease. Development 149, dev186536 (2022).

16. van Arensbergen, J., van Steensel, B. & Bussemaker, H. J. In search of the determinants of enhancer–promoter interaction specificity. Trends in Cell Biology 24, 695–702 (2014).

17. Venken, K. J. T., He, Y., Hoskins, R. A. & Bellen, H. J. P[acman]: a BAC transgenic platform for targeted insertion of large DNA fragments in D. melanogaster. Science 314, 1747–1751 (2006).

18. Fukaya, T., Lim, B. & Levine, M. Enhancer Control of Transcriptional Bursting. Cell 166, 358–368 (2016).

19. Perry, M. W., Boettiger, A. N., Bothma, J. P. & Levine, M. Shadow Enhancers Foster Robustness of Drosophila Gastrulation. Current Biology 20, 1562–1567 (2010).

20. Zeitlinger, J. et al. Whole-genome ChIP–chip analysis of Dorsal, Twist, and Snail suggests integration of diverse patterning processes in the Drosophila embryo. Genes Dev. 21, 385– 390 (2007).

21. Estrada, J., Ruiz-Herrero, T., Scholes, C., Wunderlich, Z. & DePace, A. H. SiteOut: An Online Tool to Design Binding Site-Free DNA Sequences. PLOS ONE 11, e0151740 (2016).

22. Scholes, C., Biette, K. M., Harden, T. T. & DePace, A. H. Signal Integration by Shadow Enhancers and Enhancer Duplications Varies across the Drosophila Embryo. Cell Rep 26, 2407–2418.e5 (2019).

23. Bertrand, E. et al. Localization of ASH1 mRNA Particles in Living Yeast. Molecular Cell 2, 437–445 (1998).

24. Garcia, H. G., Tikhonov, M., Lin, A. & Gregor, T. Quantitative imaging of transcription in living Drosophila embryos links polymerase activity to patterning. Curr Biol 23, 2140–2145 (2013).

25. Vera, M., Tutucci, E. & Singer, R. H. Imaging Single mRNA Molecules in Mammalian Cells Using an Optimized MS2-MCP System. in Imaging Gene Expression: Methods and Protocols (ed. Shav-Tal, Y.) 3–20 (Springer, New York, NY, 2019). doi:10.1007/978-1-4939-9674-2_1.

26. Yokoshi, M., Segawa, K. & Fukaya, T. Visualizing the Role of Boundary Elements in Enhancer-Promoter Communication. Molecular Cell 78, 224–235.e5 (2020).

27. Thomas, H. F. et al. Enhancer cooperativity can compensate for loss of activity over large genomic distances. Mol Cell 85, 362–375.e9 (2025).

28. Bénichou, O., Guérin, T. & Voituriez, R. Mean first-passage times in confined media: from Markovian to non-Markovian processes. J. Phys. A: Math. Theor. 48, 163001 (2015).

29. Keizer, V. I. P. et al. Live-cell micromanipulation of a genomic locus reveals interphase chromatin mechanics. Science 377, 489–495 (2022).

30. Marshall, W. F. et al. Interphase chromosomes undergo constrained diffusional motion in living cells. Curr Biol 7, 930–939 (1997).

31. Nozaki, T. et al. Flexible and dynamic nucleosome fiber in living mammalian cells. Nucleus 4, 349–356 (2013).

32. Yang, J. H. & Hansen, A. S. Enhancer selectivity in space and time: from enhancer– promoter interactions to promoter activation. Nat Rev Mol Cell Biol 25, 574–591 (2024).

33. Small, S., Blair, A. & Levine, M. Regulation of even-skipped stripe 2 in the Drosophila embryo. EMBO J 11, 4047–4057 (1992).

34. Frasch, M. & Levine, M. Complementary patterns of even-skipped and fushi tarazu expression involve their differential regulation by a common set of segmentation genes in Drosophila. Genes Dev 1, 981–995 (1987).

35. Macdonald, P. M., Ingham, P. & Struhl, G. Isolation, structure, and expression of *even-skipped*: A second pair-rule gene of Drosophila containing a homeo box. Cell 47, 721–734 (1986).

36. Syed, S., Duan, Y. & Lim, B. Modulation of protein-DNA binding reveals mechanisms of spatiotemporal gene control in early Drosophila embryos. eLife 12, e85997 (2023).

37. Tunnacliffe, E. & Chubb, J. R. What Is a Transcriptional Burst? Trends in Genetics 36, 288– 297 (2020).

38. Wang, Y., Ni, T., Wang, W. & Liu, F. Gene transcription in bursting: a unified mode for realizing accuracy and stochasticity. Biol Rev Camb Philos Soc 94, 248–258 (2019).

39. Berrocal, A., Lammers, N. C., Garcia, H. G. & Eisen, M. B. Kinetic sculpting of the seven stripes of the Drosophila even-skipped gene. eLife 9, e61635 (2020).

40. Ko, M. S. H. A stochastic model for gene induction. Journal of Theoretical Biology 153, 181– 194 (1991).

41. Peccoud, J. & Ycart, B. Markovian Modeling of Gene-Product Synthesis. Theoretical Population Biology 48, 222–234 (1995).

42. Singh, A., Vargas, C. A. & Karmakar, R. Stochastic analysis and inference of a two-state genetic promoter model. in 2013 American Control Conference 4563–4568 (2013). doi:10.1109/ACC.2013.6580542.

43. Leyes Porello, E. A., Trudeau, R. T. & Lim, B. Transcriptional bursting: stochasticity in deterministic development. Development 150, dev201546 (2023).

44. Nieto, C., Vahdat, Z., Lim, B. & Singh, A. Regulation of Transcriptional Bursting and Spatial Patterning in Early Drosophila Embryo Development. 2025.05.02.651973 Preprint at 10.1101/2025.05.02.651973 (2025).

45. Cho, C.-Y. & O’Farrell, P. H. Stepwise modifications of transcriptional hubs link pioneer factor activity to a burst of transcription. Nat Commun 14, 4848 (2023).

46. Di Giammartino, D. C., Polyzos, A. & Apostolou, E. Transcription factors: building hubs in the 3D space. Cell Cycle 19, 2395–2410 (2020).

47. Mir, M. et al. Dense Bicoid hubs accentuate binding along the morphogen gradient. Genes Dev 31, 1784–1794 (2017).

48. Donovan, B. T. & Larson, D. R. Regulating gene expression through control of transcription factor multivalent interactions. Molecular Cell 82, 1974–1975 (2022).

49. Trojanowski, J. et al. Transcription activation is enhanced by multivalent interactions independent of phase separation. Molecular Cell 82, 1878–1893.e10 (2022).

50. Fallacaro, S. et al. A fine kinetic balance of interactions directs transcription factor hubs to genes. bioRxiv 2024.04.16.589811 (2024) doi:10.1101/2024.04.16.589811.

51. Fallacaro, S., Mukherjee, A., Turner, M. A., Garcia, H. G. & Mir, M. Transcription factor hubs exhibit gene-specific properties that tune expression. 2025.04.07.647578 Preprint at 10.1101/2025.04.07.647578 (2025).

52. Tschopp, P., Tarchini, B., Spitz, F., Zakany, J. & Duboule, D. Uncoupling Time and Space in the Collinear Regulation of Hox Genes. PLoS Genet 5, e1000398 (2009).

53. Berrocal, A., Lammers, N. C., Garcia, H. G. & Eisen, M. B. Unified bursting strategies in ectopic and endogenous even-skipped expression patterns. eLife 12, (2024).

54. Henikoff, S. & Levens, D. L. The plusses and minuses of DNA torsion. eLife 14, e106351 (2025).

55. Liu, L. F. & Wang, J. C. Supercoiling of the DNA template during transcription. Proceedings of the National Academy of Sciences 84, 7024–7027 (1987).

56. Teves, S. S. & Henikoff, S. DNA torsion as a feedback mediator of transcription and chromatin dynamics. Nucleus 5, 211–218 (2014).

57. Chen, B.-C. et al. Lattice light-sheet microscopy: imaging molecules to embryos at high spatiotemporal resolution. Science 346, 1257998 (2014).

58. Liu, T.-L. et al. Observing the cell in its native state: Imaging subcellular dynamics in multicellular organisms. Science 360, eaaq1392 (2018).

59. Ruan, X. et al. Image processing tools for petabyte-scale light sheet microscopy data. Nat Methods 21, 2342–2352 (2024).

60. Pachitariu, M. & Stringer, C. Cellpose 2.0: how to train your own model. Nat Methods 19, 1634–1641 (2022).

61. Archit, A. et al. Segment Anything for Microscopy. Nat Methods 22, 579–591 (2025).

