## Supplementary Figures for "Enhancer Placement Impacts Transcriptional Dynamics in *Drosophila* Embryos"

### Supplemental Figures

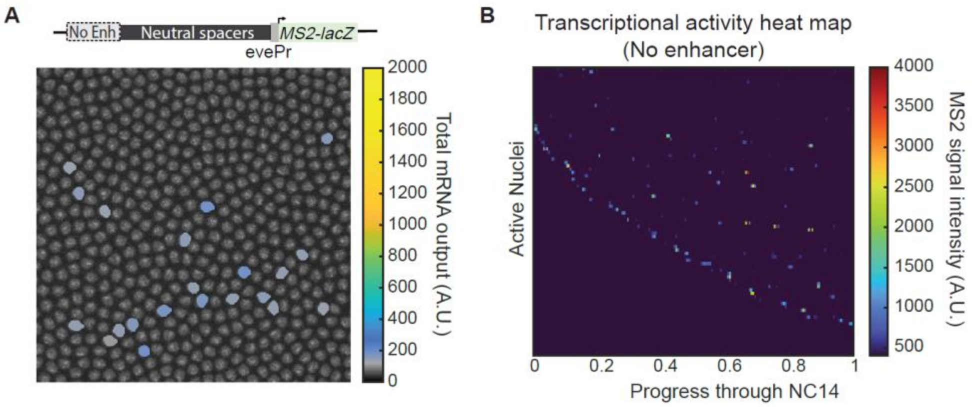


**Supplemental Figure 1. Control construct without enhancer shows negligible transcription with neutral spacers, MS2, and lacZ**

(A) Schematic of control construct lacking an enhancer sequence along with a false colored embryo snapshot depicting total mRNA produced by each nucleus in the pictured embryo. (B) Heat map of transcriptional trajectories for all active nuclei in this construct.

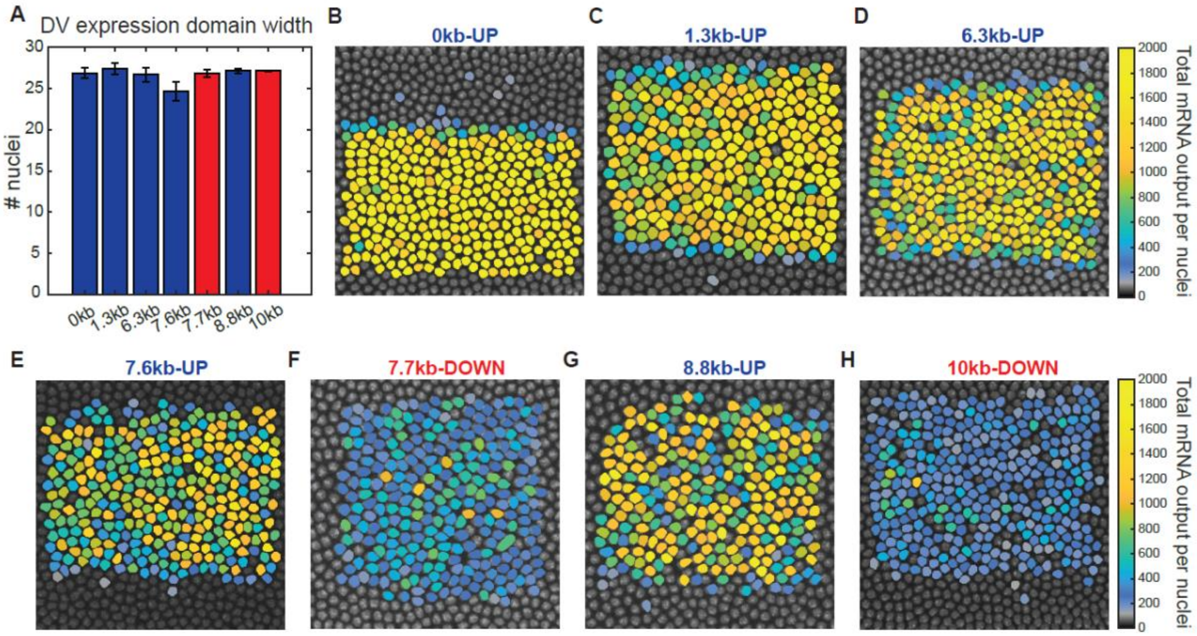


**Supplemental Figure 2. Enhancer-promoter configuration does not significantly affect expression domain or activity pattern**

(A) Width of dorsoventral expression domain driven by each *MS2-lacZ* reporter under varied enhancer-promoter configurations, reported as number of nuclei. Each data set contains combined data for 3 biological replicates. Error bar represents standard error of the mean. (B-H) False colored embryo snapshot depicting total mRNA produced by each nucleus.

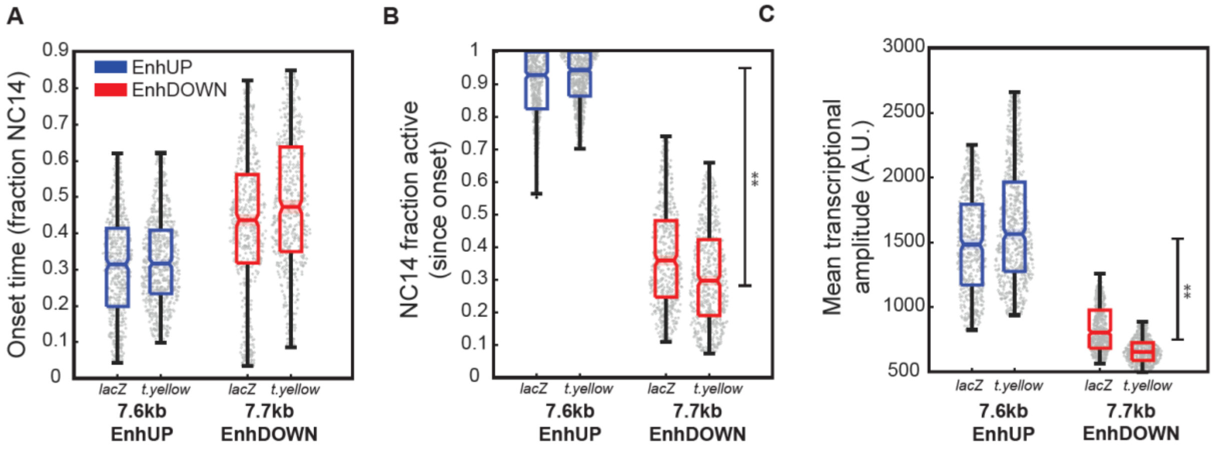


**Supplemental Figure 3. Metrics of transcription kinetics are not significantly affected by the replacement of *lacZ* with the *t.yellow* reporter in mid-range constructs with opposing enhancer position**

(A) Onset time plotted for 7.6 kb-EnhUP and 7.7 kb-EnhDOWN constructs with either *lacZ* or *t.yellow* reporter genes. (B) Fraction of NC14 after onset time that each nucleus spends in the active state plotted for each shown configuration. (C) Mean transcriptional amplitude of the active state plotted for each shown configuration. Statistical significance between EnhUP and EnhDOWN subsets in B and C, performed by combining the data into two groups based on position and analyzing by student's t-test with α=0.05. Each data set contains combined data for 3 biological replicates. Number of nuclei in each data set as follows: n(7.6kb-*lacZ*)=991, n(7.6kb-*t.yellow*)=909, n(7.7kb-*lacZ*)=681, n(7.7kb-*t.yellow*)=801.

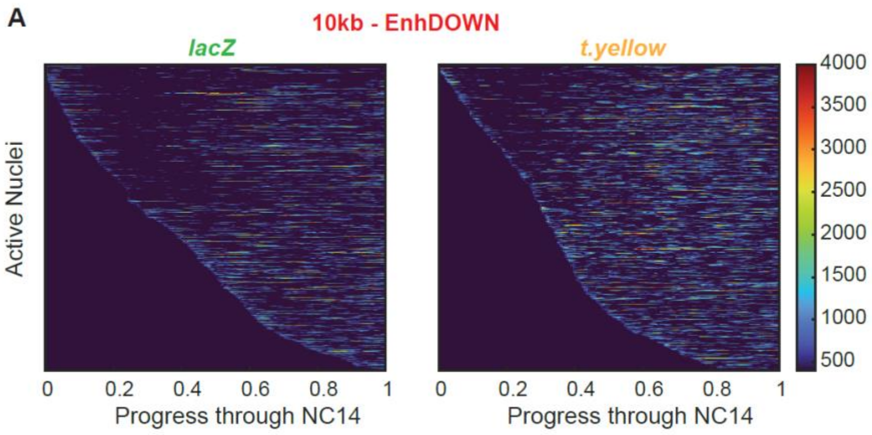


**Supplemental Figure 4. Transcription kinetics are not significantly affected by the replacement of *lacZ* with the *t.yellow* reporter 10 kb construct**

(A) Heat map of transcriptional trajectories for all active nuclei in the 10 kb-EnhDOWN constructs with either *lacZ* (left) or *t.yellow* (right) reporter genes.
